# Genome sequence of the medicinal plant *Urtica dioica* reveals the genetic basis of the flavonoid metabolism

**DOI:** 10.64898/2026.05.15.725508

**Authors:** Katharina Wolff, Julie Anne V.S. de Oliveira, Lena Fürstenberg, Marie Hagedorn, Bennet Garz, Milan Borchert, Boas Pucker

## Abstract

**Background:** *Urtica dioica*, also known as stinging nettle, is a widespread plant that can indicate high nitrogen availability in the soil. It is probably best known for the pain caused by touching it. *U. dioica* is also recognized as a medicinal plant with reports claiming applicability against numerous diseases.

**Results:** A highly continuous genome sequence was constructed based on nanopore long read sequencing data. The total assembly size is 1.1 Gbp with an N50 of 40.7 Mbp. RNA-seq data and hints from other species were integrated to produce a high quality annotation of the protein encoding genes. This genomic resource enabled the identification of genes involved in the flavonoid biosynthesis. A particular focus was on anthocyanin biosynthesis genes as these are crucial for high light and nitrogen deprivation stress response, which is revealed by redding of the leaves.

**Conclusion:** This genomic resource provides the basis for future studies unraveling the biosynthesis pathways underlying various medically important compounds produced by stinging nettles.

## Background

*Urtica dioica* (**Fig. 1**) or commonly known as stinging nettle is a widespread plant whose presence indicates high nitrogen availability in the soil (Li, 1994). Due to *U. dioica’s* ability to form rhizomes, it is able to grow perennially, allowing it to thrive in various environments with moderate climate and sufficient rain (Taylor, 2009). Therefore, *U. dioica* can be found in different parts of the world, such as Europe, Asia, North America and North Africa and is commonly regarded as a weed due to its fast growth.

**Fig. 1:**
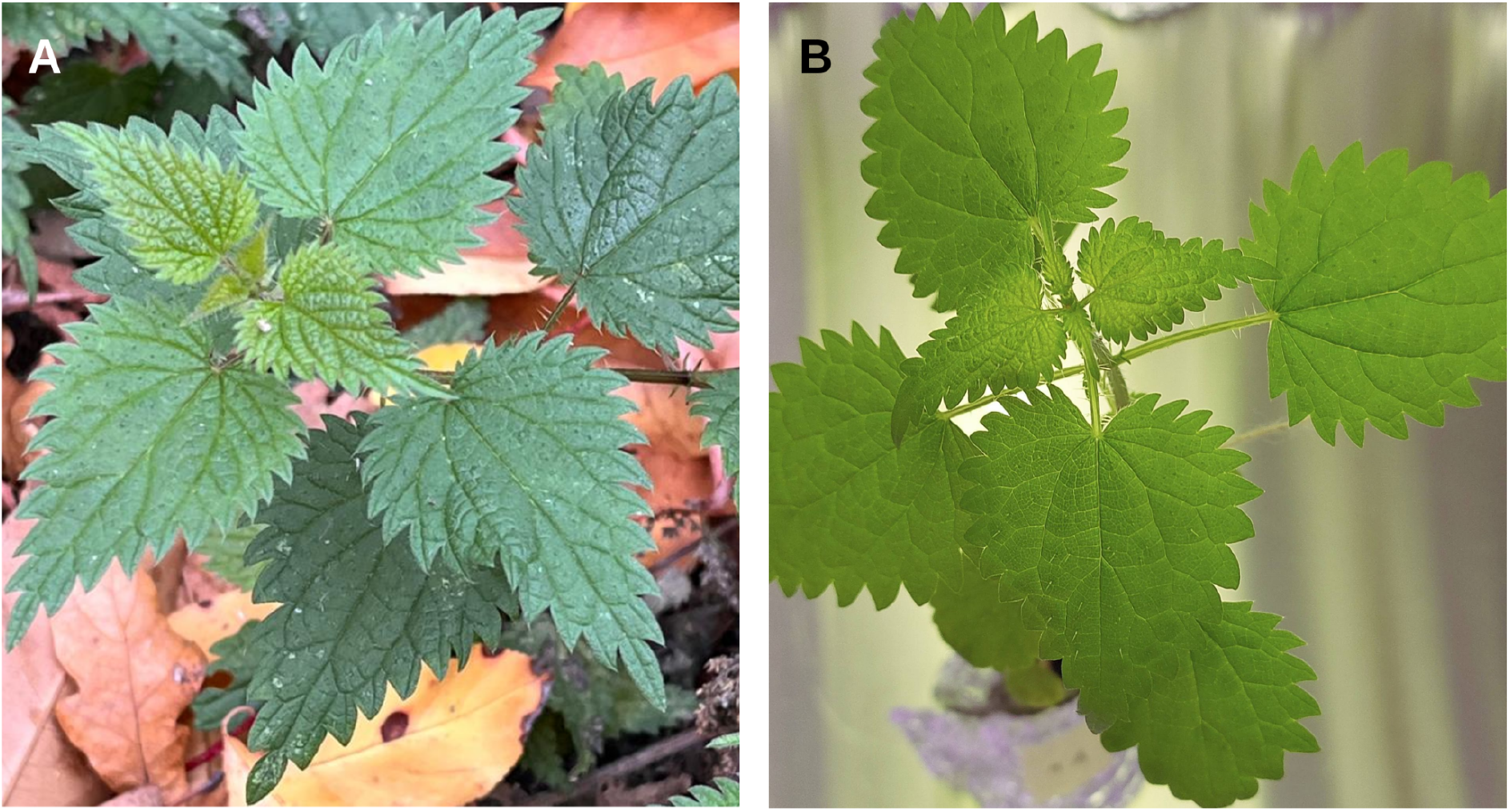
Picture of *Urtica dioica*. Plant growing in the wild (A) and plant cultivated in the lab. Photo credit: Katharina Wolff and Lena Fürstenberg.

Stinging nettle is probably best known for the pain experienced when touching its leaves, which is attributed to hollow hairs (trichomes) that act as hypodermic needles causing mechanical and biochemical damage (Thurston & Lersten, 1969; Cummings & Olsen, 2011; Ensikat *et al*., 2021). A chemical cocktail involving inflammatory compounds is injected from a gland at the base of the hair into the skin when a sealing bulbous tip is broken off the hairs at contact (Anderson *et al*., 2003; Cummings & Olsen, 2011). It is assumed that histamine, acetylcholine, and serotonin play an important role in this chemical defense of stinging nettles, but the observed effects cannot be fully explained by the compounds identified yet, suggesting that an additional neurotoxin might be involved (Emmelin & Feldberg, 1947; Schildknecht, 1981; Tuberville *et al*., 1996; Ensikat *et al*., 2021). The main ecological function of stinging hairs is assumed to be defense against mammalian herbivores (Ensikat *et al*., 2021).

*Urtica dioica* has been utilized in traditional medicine for many decades and is known for both its nutritional and medicinal value (Baumgardner, 2016). The use of medicinal plants has a long standing history and played an important role in the promotion of human health. It is important to emphasize that the use of a plant in traditional medicine might indicate the effectiveness against a certain illness, but additional evidence is needed. Nevertheless, traditional knowledge can lead to studies identifying active compounds in plants that might inspire the development of drugs (Gupta *et al*., 2005; Dutfield, 2010; Pirintsos *et al*., 2022). Generally, medicinal plants are characterized by a high amount of bioactive specialized secondary metabolites. Identifying the genes for the production of specialized plant compounds has developed into an intensely studied field within the life sciences (Kang *et al*., 2020; Polturak *et al*., 2022; Siadjeu & Pucker, 2023). *U. dioica* was celebrated as the medicinal plant of the year 2022 in Germany. A range of phytomedical compounds have been reported in *U. dioica* including phenolic compounds and sterols (Alimoddin *et al*., 2024). Several phytocompounds occurring in *U. dioica* have been reported to exhibit health promoting benefits such as anti-diabetic, anti-rheumatic, anti-microbial and anti-asthmatic effects during internal application (Joshi *et al*., 2014; Esposito *et al*., 2019; Abd-Nikfarjam *et al*., 2022). For example, stinging nettle tea is frequently consumed due to the reported health benefits. *U*.*dioica* has been reported to contain a high amount of a structurally and functionally diverse group of specialized metabolites the flavonoids (Esposito *et al*., 2019; Tarasevičienė *et al*., 2023; Koczkodaj *et al*., 2023; Semwal *et al*., 2023). Flavonoids represent a versatile group of specialized plant metabolites, which consists of several subgroups, including: flavonols, flavones, anthocyanins, and proanthocyanidins and are most commonly associated with their strong antioxidant activity (Winkel-Shirley, 2001). Due to this characteristic, flavonoids are involved in stress response within the plant. The formation of flavonoids is associated with a broad range of stressors such as drought, heat, cold, ultraviolet radiation, high light intensities and specific ion concentrations in the soil (Pucker & Selmar, 2022; Grünig *et al*., 2025). The fundamental biosynthesis of flavonoids is highly conserved across species and represents one of the most studied specialized biosynthesis pathways (Winkel-Shirley, 2001; Grotewold, 2006; Grünig *et al*., 2025).

While *Urtica dioica* has a long tradition of use as a medicinal plant and produces a variety of bioactive compounds, the genetic basis of bioactive compounds biosynthesis has remained largely unexplored. In this study, a genome sequence and corresponding annotation of *U. dioica* are provided and utilized to explore the genetic mechanism for the production of flavonoids, focusing on anthocyanins.

## Results

### Genome sequence assembly and annotation

A total of about 53.48 Gbp of ONT R9 sequencing data was generated, consisting of 2,192,237 reads with a raw read N50 of 44.11 kbp (Additional file A). Additionally, 73.17 Gbp of ONT R10 sequencing data was generated, consisting of 7,513,588 reads with a raw read N50 of 39.83 kbp (Additional file A). Different assembly approaches were conducted with R9 and R10 datasets assessing the performance of different assemblers to obtain the best genome sequence (Additional file C).

The selected genome sequence was generated by Hifiasm and has a size of 1.1 Gbp (**Table 1**) with an N50 of 40.67 Mbp. This genome sequence is available via bonndata (10.60507/FK2/1XHSZ3). While this assembly size is substantially smaller than the previous estimate, this might be due to methodological and biological differences. A high completeness of the genome sequence was indicated by the detection of 97.5% of complete BUSCO genes(C:97.5%[S:16.7%,D:80.8%],F:1.3%,M:1.1%,n:2026,E:2.0%; based on embryophyta_odb12). The high percentage of duplicated BUSCO genes indicates that haplophases have been separated.

**Table 1:** Summary of selected assembly statistics and assessment of completeness based on BUSCO score. The different assemblers are listed followed by the data type that went into each assembly (R9 or R10 flow cells and corresponding sequencing chemistry).

|  | Shasta (R9) | NextDenovo2 (R9) | Hifiasm (R10) |
| --- | --- | --- | --- |
| **number of contigs:** | 7,562 | 1,464 | 491 |
| **maximal contig length** | 9.1 Mbp | 11.6 Mbp | 60.9 Mbp |
| **total number of bases** | 1 Gbp | 1.36 Gbp | 1.1 Gbp |
| **GC content:** | 41% | 41% | 42.3% |
| **N50** | 0.33 Mbp | 1.42 Mbp | 40.67 Mbp |
| **N90** | 0.07 Mbp | 0.47 Mbp | 1.27 Mbp |
| **BUSCO (complete)** | 93.7% | 96.8% | 97.5% |

The gene prediction with GeMoMa revealed a total of 37456 protein encoding genes. The identification of 96.9% of complete BUSCO genes in a set of representative polypeptide sequences, in which each gene was only represented by one sequence, indicates high completeness. This value is only slightly lower than the 97.5% of completeness observed for the genome sequence, which indicates that the structural annotation process has captured most genes.

The genome sequence completeness and separation of haplophases was assessed based on coverage analyses. A long-read mapping against the genome sequence revealed the overall coverage distribution showing peaks at approximately 25x and 50x resulting in an average coverage depth of around 40x (**Fig. 2A**). The MGSE (Natarajan *et al*., 2025) concept was applied to infer a genome size of about 1.3 Gbp based on the total amount of sequencing data of 53.48 Gbp. A frequency analysis of the “duplicated” BUSCO genes showed that two copies is the dominant situation thus suggesting that many gene loci are represented by contigs of two haplophases (**Fig. 2B**).

**Fig. 2:**
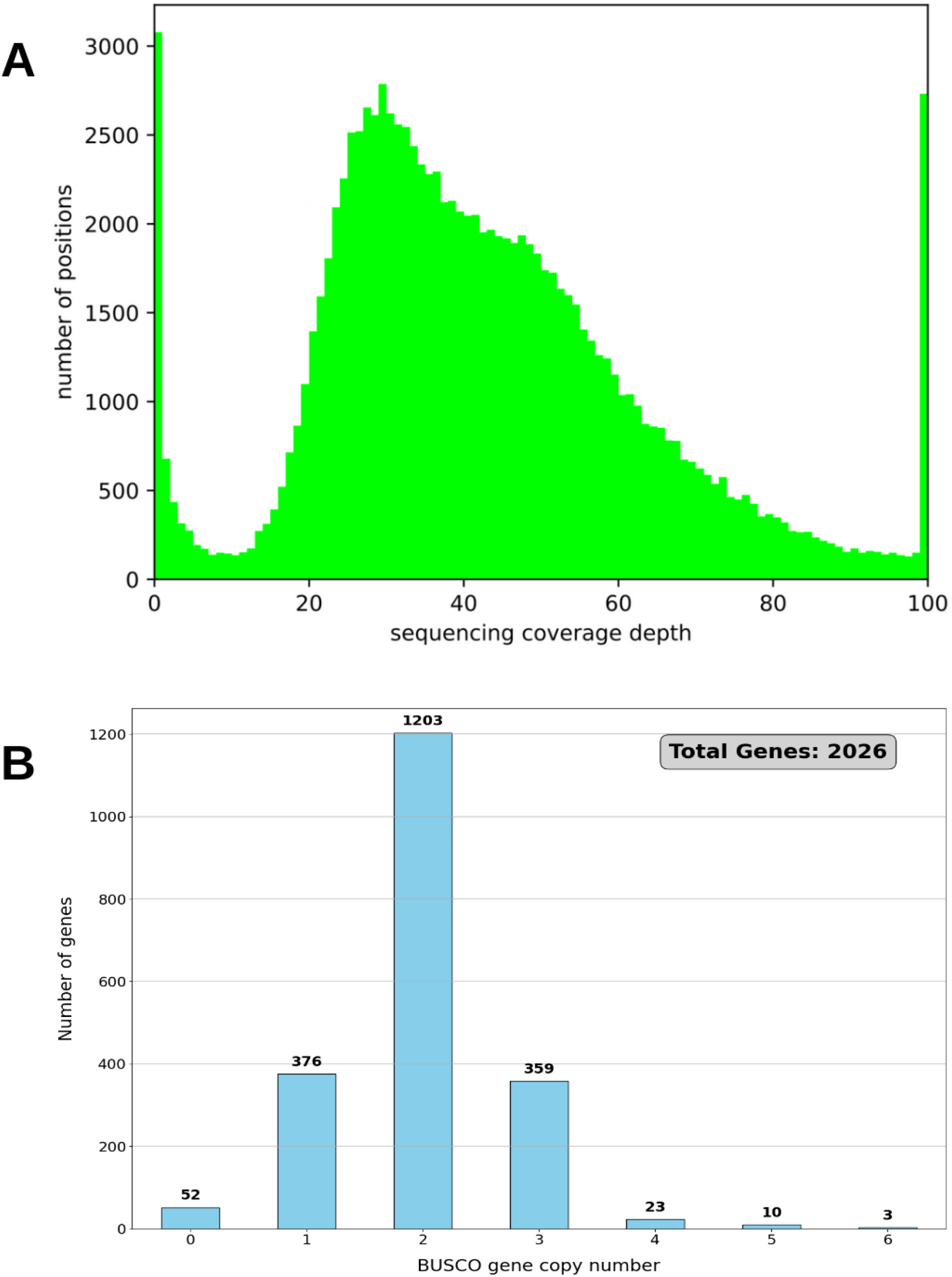
Coverage of a long-read mapping of all R9 sequencing data to the *Urtica dioica* genome sequence (A) and frequency analysis of “duplicated” BUSCO gene copy numbers (B).

### Identification of flavonoid biosynthesis genes in *Urtica dioica*

One objective of this study was to identify candidate genes which play a role in the flavonoid biosynthesis in *U. dioica*. Candidate genes encoding enzymes for all steps in the anthocyanin and proanthocyanidin biosynthesis were identified via KIPEs (**Fig. 3**). While most enzymatic steps of the flavonoid biosynthesis are covered by candidates displaying all known functionally important amino acid residues in the correct positions, some members of the 2-oxoglutarate-dependent dioxygenase (2-OGDDs) family show deviations. This concerns the flavonol synthase (FLS), the first committed step of the flavonol biosynthesis branch, and the anthocyanidin synthase (ANS), a crucial step in the anthocyanin biosynthesis. Further investigations are needed to determine whether these are lineage-specific modifications, which might alter the enzyme properties. Transcription factors of the MYB and bHLH families involved in the activation of the anthocyanin biosynthesis through the MBW complex were identified, too. This includes MYB75/PAP1 for the activation of the anthocyanin biosynthesis and MYB123/TT2 for the activation of the proanthocyanidin biosynthesis.

**Fig. 3:**
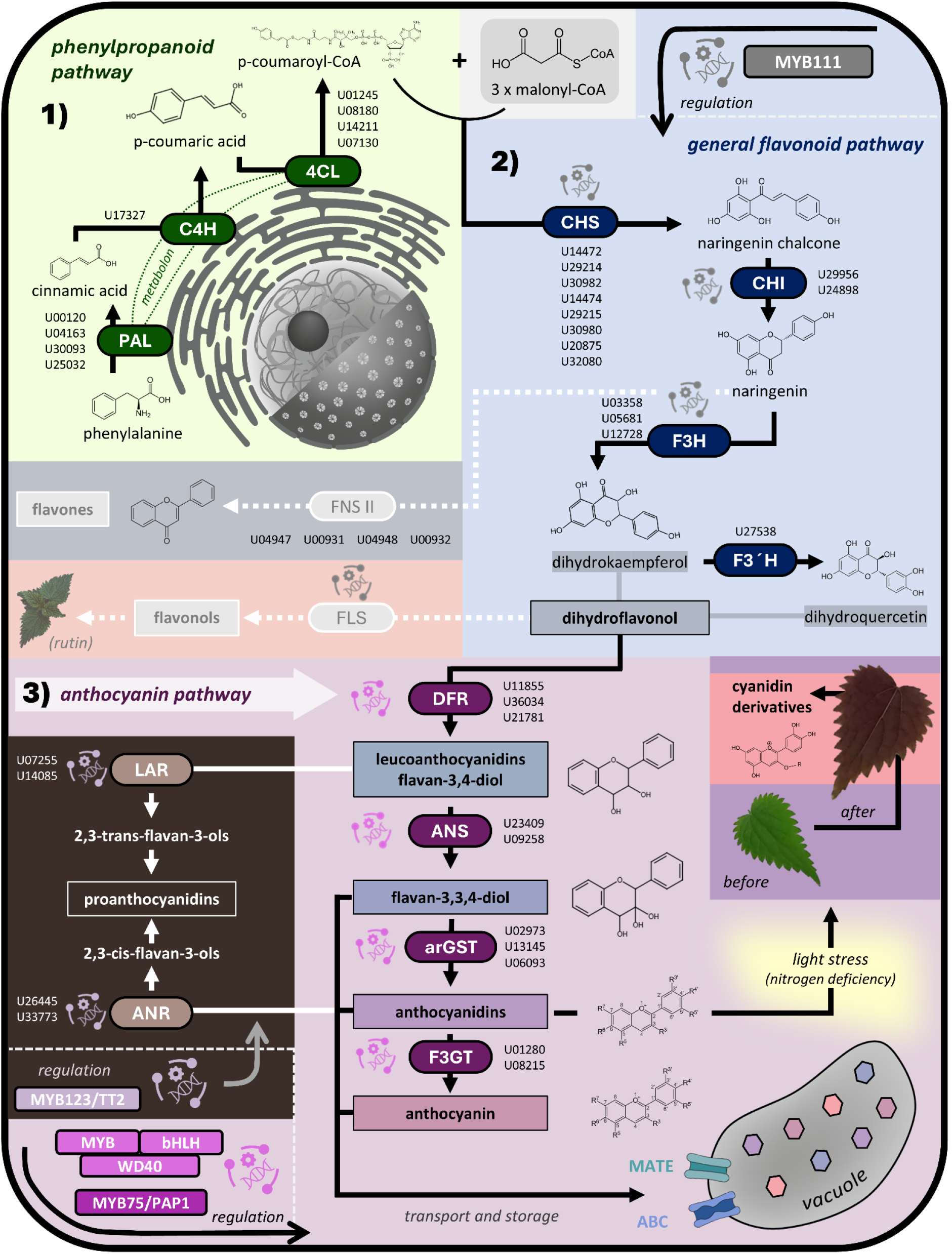
Simplified representation of the flavonoid biosynthesis pathway in *Urtica dioica* with indicated candidate genes. The following pathways are represented: 1) phenylpropanoid pathway, 2) general flavonoid pathway and 3) anthocyanin pathway. The genes involved are abbreviated as follows: PAL - phenylalanine ammonia-lyase, C4H - cinnamic acid 4-hydroxylase, 4CL−4-coumarate-CoA ligase, CHS - chalcone synthase, CHI - chalcone isomerase, F3H - flavanone 3-hydroxylase, F3’H - flavonoid 3’-hydroxylase, FNS II - flavone synthase II, FLS - flavonol synthase, DFR - dihydroflavonol 4-reductase, ANS - anthocyanidin synthase, arGST – anthocyanin-related glutathione S-transferase, 3GT - anthocyanidin-3-O-glucosyltransferase, LAR - leucoanthocyanidin reductase and ANR - anthocyanidin reductase. The displayed gene IDs have been shortened by replacing “Udioica” with “U”.

### Expression of flavonoid biosynthesis genes

Expression of the flavonoid biosynthesis genes was assessed across a variety of different plant structures (**Fig. 4**). Candidate genes showed expression in at least one tissue, which supports their validity. High activity of genes involved in the proanthocyanidin biosynthesis was observed in the roots. The anthocyanin biosynthesis does not show strong activity in any of the investigated tissues, but the core genes of the anthocyanidin biosynthesis branch, namely DFR, ANS, arGST, are active in fibres.

**Fig. 4:**
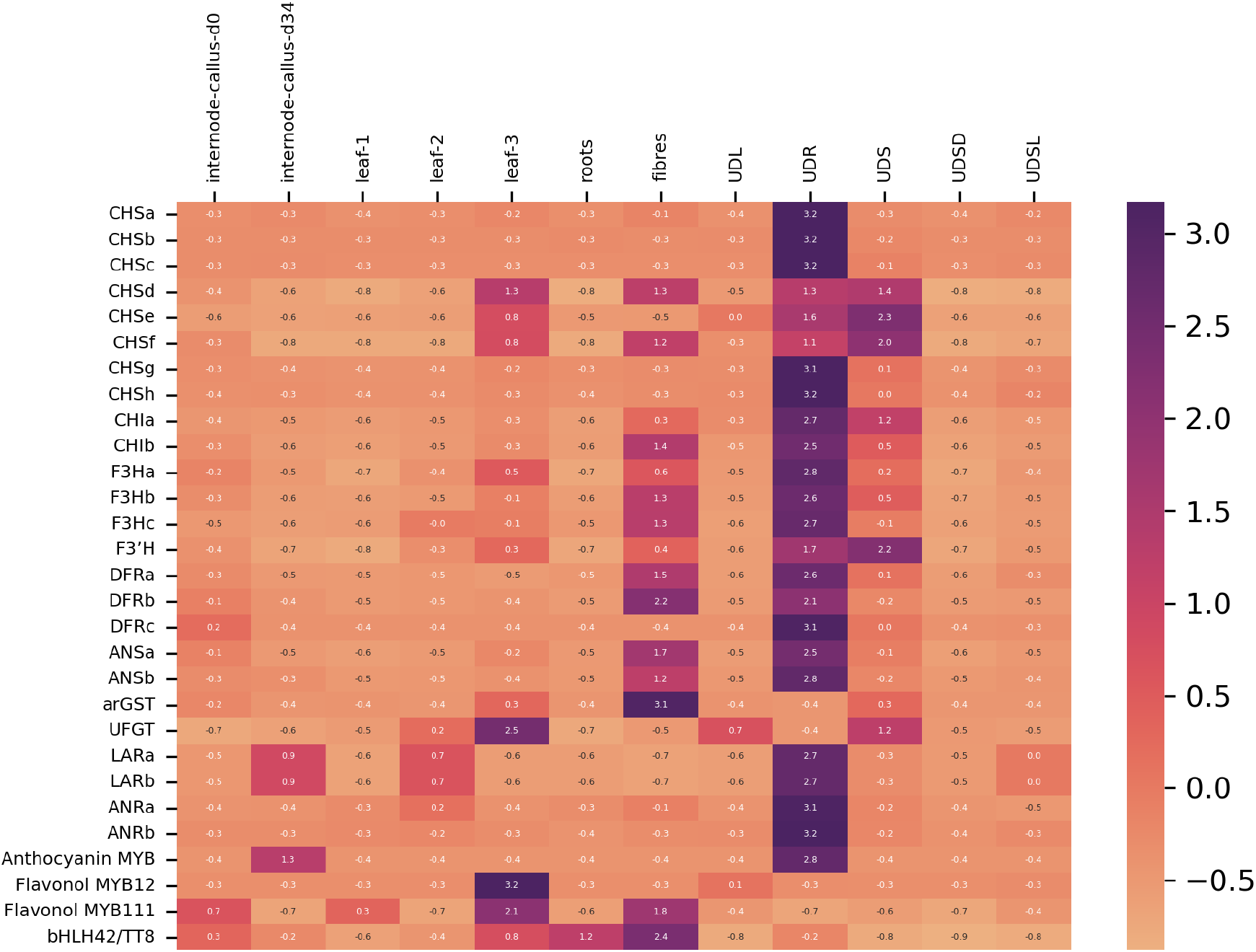
Gene expression across different *Urtica dioica* tissues. Displayed genes encode chalcone synthase (CHS), chalcone isomerase (CHI), flavanone 3-hydroxylase (F3H), flavonoid 3’-hydroxylase (F3’H), dihydroflavonol 4-reductase (DFR), anthocyanidin synthase (ANS), anthocyanin-related glutathione S-transferase (arGST), leucoanthocyanidin reductase (LAR), anthocyanidin reductase (ANR), MYB75, MYB123, MYB111/12, TRANSPARENT TESTA 8 (TT8). Only active copies of these genes are displayed here, while a full heatmap is available as Additional file B. Samples: UDL (leaf), UDS (stem), UDSD (seed), UDSL (seedling), and UDR (roots). Displayed expression values have been z-score normalized.

### Stress induced anthocyanin production in *U. dioica*

The impact of high light intensity (HL) and nitrogen starvation (ND) on the activation of the anthocyanin biosynthesis in *U. dioica* was explored in a stress experiment. At the start of this experiment, all plants displayed green leaves with negligible anthocyanin pigmentation. Two weeks of environmental stress caused the leaves of the HL-treated plants to exhibit a notable color change towards purple, with anthocyanin content increasing by approximately 8-fold compared to the control group (**Fig. 5**). The combination of HL+ND showed an even stronger increase, with an 11-fold rise in anthocyanin levels. In contrast, the ND group showed only a slight increase in anthocyanin content, about twofold. The control group remained largely unchanged over the time of this experiment.

**Fig. 5:**
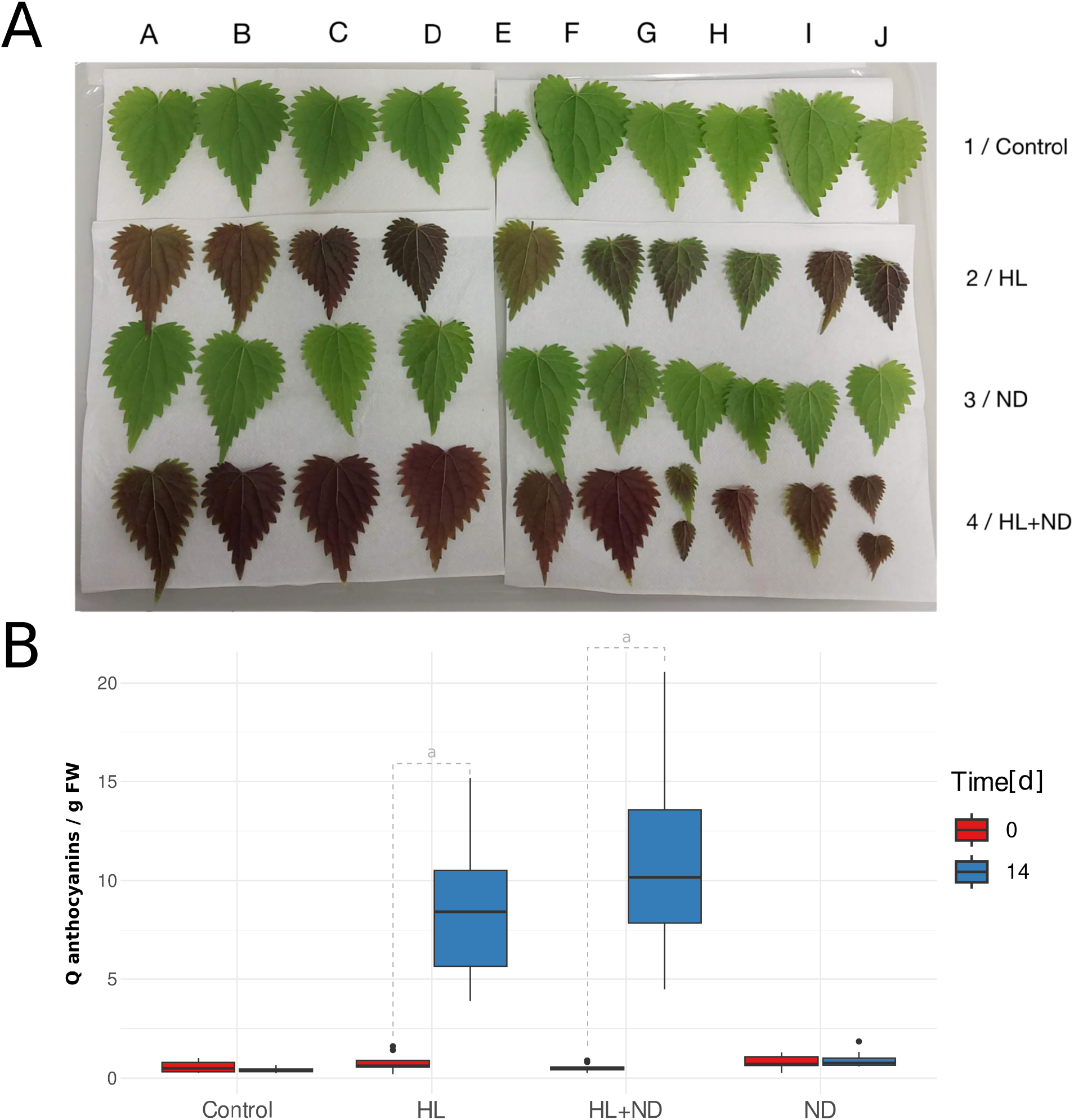
**A:** Pictures of plant leaves exposed to different treatments over two weeks. Replicates (A-J) show diversity in leaf morphology. Control plants were grown under normal light (20-50 µmol m^-2^ s^-1^ PPFD), high light (HL, 180-200 µmol m^-2^ s^-1^ PPFD) was applied for two weeks, nitrogen deficiency (ND) was achieved with Hoagland solution without nitrogen. **B:** Boxplot (median and interquartile range) visualizing the relative anthocyanin content (Q) per gram FW of plant samples grown under different stress conditions (n = 10) at day 0 and day 14 of growth. The conditions are annotated as HL: high light stress, ND: Nitrogen deficiency, HL+ND: High light stress and nitrogen deficiency. “a” indicates a significant difference within the group with a p value < 0.001. The raw values can be found in Additional file F.

It was also observed that younger leaves in the HL and HL+ND groups exhibited stronger purple pigmentation compared to older leaves, which were more red in color. This color change was particularly noticeable on the abaxial side of the leaves, while the adaxial side showed a darker green with a slight purple tint.

An additional control was conducted by covering leaves partially with aluminum foil, which blocked light exposure (Additional file E). The covered leaf sections remained green, while the light exposed areas of the same leaf turned red or purple, suggesting that anthocyanins accumulate specifically in areas exposed to high light.

## Discussion

### Genome sequence of *Urtica dioica* reveals flavonoid biosynthesis genes

The genome sequence generated as part of this study provides a solid basis for access to the gene repertoire of *Urtica dioica*. The assembly size of 1.1 Gbp in our study indicates that two haplophases are represented given the size of 618 Mbp previously reported for one haplophase of this species (Christenhusz *et al*., 2025). The BUSCO gene copy analysis supports the assumption that the plant analyzed in this study is diploid with a major peak showing two copies. In general, the completeness of the genome sequence and annotation appears to exceed those of previously published genome sequences (Christenhusz & Twyford, 2024; Christenhusz *et al*., 2025; Hirabayashi *et al*., 2026). The continuity of this *U. dioica* genome sequence is similar to scaffold level continuity assemblies of *U. urens* (Christenhusz & Twyford, 2024) and *U. dioica* (Hirabayashi *et al*., 2026). Given the high quality of this genome sequence, it is surprising that no suitable candidate gene for a flavonol synthase was discovered. It is even more striking, because *U. dioica* is well known for the production of flavonols, especially rutin (Chaurasia & Wichtl, 1987; Pinelli *et al*., 2008; Vajić *et al*., 2015; Koraqi *et al*., 2023).

The gene expression pattern suggests activity of the proanthocyanindin biosynthesis in the roots of stinging nettle (**Fig. 4**). As expected, no green structure of stinging nettle is showing active anthocyanin biosynthesis. This indicates that anthocyanin biosynthesis is only activated when required i.e. under stress conditions.

### Stress experiments

The results of this study confirm the hypothesis that high light conditions can strongly induce anthocyanin production in *Urtica dioica*, consistent with findings in other plant species (Zhang *et al*., 2018; Cohen *et al*., 2019; Zheng *et al*., 2021; Nowak *et al*., 2024). In this work, anthocyanin concentration correlated with leaf color which was reported previously in other plants (Guo *et al*., 2019). The observed color changes from green to purple and the associated increase in anthocyanin content in the HL and HL+ND groups suggest that these pigments may serve a photoprotective function, helping to mitigate the oxidative damage caused by excess light. Anthocyanins are probably absorbing excess light, reducing the load on photosynthetic pigments, and they may also play a role in scavenging ROS (Feild *et al*., 2001; Zeng *et al*., 2010; Zheng *et al*., 2019, 2021; Grünig *et al*., 2025). Interestingly, while anthocyanin production increased significantly under HL and HL+ND treatments, nitrogen deficiency alone did not result in a substantial increase in anthocyanins. This finding contrasts with previous studies, which suggested that nitrogen deficiency could directly stimulate anthocyanin biosynthesis (Scheible *et al*., 2004; Liang & He, 2018). One explanation for this discrepancy is that nitrogen deficiency alone may not induce significant stress in the absence of high light exposure. It is possible that low nitrogen levels made the plants more susceptible to light-induced damage, but without the intensity of high light, the need for photoprotective anthocyanins was minimal. Additionally, nitrogen content in the leaves is reported to decrease much slower in maize plants in low light conditions (Ding *et al*., 2005). If the nitrogen content in the stinging nettle leaves did not decrease substantially in the two weeks of this experiment, no nitrogen deficiency would have occurred. This should be further investigated in future studies.

The combination of high light and nitrogen deficiency (HL+ND) resulted in the highest anthocyanin concentrations, suggesting a synergistic effect between these two stressors. Nitrogen deficiency may have impaired the plants’ ability to cope with high light, thus triggering an increased need for anthocyanins to protect the leaves from photooxidative damage. This is in line with studies that have shown a link between nitrogen availability and the plant’s ability to withstand high light stress (Peng *et al*., 2007; Liang & He, 2018; Cohen *et al*., 2019). Leaf parts covered with aluminum foil and thereby shielded from high light intensity, remained green, while the light exposed areas of the leaf turned red. This suggests that anthocyanin biosynthesis is specifically activated in the light exposed areas and that anthocyanins are produced locally and not transported to other parts of the leaf. This observation also aligns with previous observations in the flowers of *Victoria cruziana*, which showed pigmentation in areas not covered by aluminum foil (Nowak *et al*., 2024). Our findings corroborate the hypothesis that anthocyanins are only synthesized in response to environmental cues when beneficial for the plant, such as for protection against light stress. It also indicates that nitrogen deficiency may indirectly influence anthocyanin biosynthesis, as the entire leaf would likely have shown pigmentation if nitrogen deficiency alone could directly trigger anthocyanin production. In our experiments, high light exposure appeared as a much stronger trigger for anthocyanin formation than nitrogen starvation.

## Conclusions

This study reports a genome sequence with corresponding annotation for *Urtica dioica*, which enabled the identification of flavonoid biosynthesis genes. Expression of these genes was explored in a variety of tissues of this biomedical plant. The activation of the anthocyanin biosynthesis by light treatment and nitrogen deficiency was investigated and revealed light as the most important trigger of pigment accumulation in stinging nettle.

## Methods

### Plant materials

The *Urtica dioica* plant (XX-0-BONN-49798) that was used for the genome sequencing was originally collected in Braunschweig, Germany. The plant was grown from a single root in hydroponic conditions. This ensured that all shoots have the same genetic makeup.

For the DNA extraction for R9 flow cell sequencing, the plants were placed under artificial grow lights (NIELLO GS600W) for a 16 h light and 8 h dark cycle and incubated at 18-23 °C. Prior to sampling, the plants were incubated in the dark for three days to reduce the amount of starch, sugar, and chloroplasts present in the leaves. Young leaves were harvested in a nondestructive manner and used for the DNA extraction.

For the DNA extraction for R10 flow cell sequencing, the plants were grown in the University of Bonn Botanic Gardens. Samples were taken on July 14th 2026 and October 6th 2025.

For RNA extraction, plants were placed under artificial grow lights (LED T5 36W 5050lm 830 warm white; LED T5 36W 5600lm 840 cold white) for a 16 h light and 8 h dark cycle and incubated at approximately 20°C.

### DNA extraction and nanopore sequencing

The general workflow followed a previously established protocol for plant genome sequencing (de Oliveira *et al*., 2026). The high molecular DNA extraction was conducted using a CTAB-based method according to the protocol “Plant DNA extraction and preparation for ONT sequencing” (Siadjeu *et al*., 2020). Next, a NanoDrop measurement, agarose gel electrophoresis, and Qubit fluorometer measurement were used to determine DNA quality and quantity. In order to achieve an optimal DNA fragment length distribution for long read sequencing, the Short Read Eliminator (SRE) kit (Pacific Biosciences) was employed, using 50 µL of each DNA sample. After the size selection of the extracted DNA, the sample was quantified again with a Qubit fluorometer in order to obtain the final concentration values. Next, a library preparation was carried out, following the Oxford Nanopore DNA end-prep and adapter ligation protocol SQK-LSK109 (ONT) for the R9 flow cell sequencing or SQK-LSK114 for the R10 flow cell sequencing, respectively. The sequencing was conducted on MinIONs and a PromethION 2 Solo according to the standard ONT sequencing protocol.

Basecalling was conducted on a GPU in the de.NBI cloud using dorado (Oxford Nanopore Technologies). For R9 data, the dna_r9.4.1_e8_hac@v3.3 model was used with dorado v0.9.6. For R10 data, HAC basecalling considering base modifications (--modified-bases 5mCG_5hmCG) was conducted with dorado v1.4.0. HERRO (Stanojević *et al*., 2026) correction of the R9 data was performed with dorado v1.1.0 and error correction of the R10 data was performed with dorado v.1.2.0. A quality control of the resulting data was executed using the Python script FASTQ_stats3.py (de Oliveira *et al*., 2026).

### Assembly

Sequencing runs were concatenated by flow cell type into a single file for R9 and R10, respectively. Assemblies were performed using different long read assemblers including NextDenovo2 (Hu *et al*., 2024), Shasta (Shafin *et al*., 2020), and Hifiasm (Cheng *et al*., 2021). Shasta v.0.10.0 was run using the default settings contained in the provided configuration file Nanopore-May2022.conf with a minimum read length of 10 kbp. NextDenovo v2.5.2 (Hu *et al*., 2024) was run with the default parameters for raw ONT reads, with a minimal read length of 10 kbp and an expected genome size of 1.6 Gbp. Hifiasm (Cheng *et al*., 2021) v0.25.0-r726 was run with all R10 reads as input.

The completeness of the resulting assemblies was assessed using BUSCO v.5.8.2 using embryophyta_odb12 (Simão *et al*., 2015; Tegenfeldt *et al*., 2025) a tool that assesses the completeness of genome sequences based on universal single-copy genes. The long-reads were mapped back to the assembled genome sequence with minimap2 v2.27-r1193 (Li, 2018). The resulting SAM file was converted into BAM with samtools v1.23.1 (Li *et al*., 2009; Danecek *et al*., 2021). The coverage depth per position was calculated with construct_cov_file.py v0.3 (Pucker & Brockington, 2018). Visualization of the coverage per position was achieved with cov_plot.py v0.4 (Pucker *et al*., 2019). The copy numbers of duplicated BUSCO genes were analyzed with a previously developed Python script busco_cov_hist_2.py (Wolff *et al*., 2026).

### Structural and functional annotation

For the structural annotation, RNA-seq data sets of *U. dioica* were mapped against the genome sequence with STAR v2.7.11b (Dobin *et al*., 2013; Dobin & Gingeras, 2015). A variety of plant tissues have been sampled for the generation of RNA-seq data including leaf (ERR17204399), stem (ERR17204400), seed (ERR17204401), seedling (ERR17204402), and roots (ERR17204403). The structural annotation was generated with GeMoMA v1.9 (Keilwagen *et al*., 2016, 2019). Genome sequences and the corresponding annotations of *Morus notabilis* (GCF_000414095.1), *Cannabis sativa* (GCF_029168945.1), *Ziziphus jujuba* (GCF_031755915.1), and *Humulus lupulus* (GCF_963169125.1) were used as references. The initial annotation was filtered with the GeMoMa Annotation Finalizer using the parameters f=“start==‘M’ and stop==‘*’ and aa>=50 and avgCov>0 and (isNaN(bestScore) or bestScore/aa>=1.5) and iAA>=0.3 and pAA>=0.3” atf=“iAA>0.9 and pAA>0.9 and avgCov>0 and tpc>=0.5 and tie>=0.5”. The completeness of both annotations was assessed by “BUSCO” v.5.8.2 (Simão *et al*., 2015; Tegenfeldt *et al*., 2025) run in protein mode with the reference dataset “embryophyta”.

Based on the structural annotation, a detailed functional annotation of the flavonoid biosynthesis genes was conducted utilizing KIPEs v3.2.7 (Rempel *et al*., 2023) with the FlavonoidBioSynBaits_v3.3, which includes bait sequences belonging to functional enzymes in other plant species. The anthocyanin biosynthesis regulating MYBs were identified with the MYB_annotator v1.0.1 (Pucker, 2022) and the anthocyanins biosynthesis regulating bHLH TT8 was identified with the bHLH_annotator v1.0.4 (Thoben & Pucker, 2023). Additionally a general functional annotation was performed with a Python script construct_anno.py (Pucker & Iorizzo, 2023), which assigns annotation terms based on the Araport11 annotation of *Arabidopsis thaliana* (Lamesch *et al*., 2012; Cheng *et al*., 2017).

### Gene expression analysis

RNA was extracted from different *U. dioica* tissues with the NucleoSpin RNA Plant and Fungi kit (Machery & Nagel) protocol and subjected to paired-end RNA-seq. These plants were grown under the same conditions as for the R10 data generation. Additionally, all *Urtica dioica* RNA-seq datasets were retrieved from the Sequence Read Archive via fastq-dump of the sra-toolkit v2.9.6.1 (NCBI, 2020; Katz *et al*., 2022). All FASTQ file pairs were processed with kallisto v0.44.0 (Bray *et al*., 2016) based on the coding sequences of the structural annotation generated in this study. Individual count tables were merged with previously developed Python scripts (Pucker & Iorizzo, 2023). A heatmap of selected genes and samples was generated with the Python script heatmap_plotter.py v0.4 (https://github.com/bpucker/PBBtools).

### Stress induced anthocyanin production in nettle

For two weeks, stinging nettle roots were kept in water under artificial grow lights (NIELLO GS600W) which emitted around 20-50 µmol/m2 s PPFD for a 16-hour photo-period at a temperature of 18-23 °C. Emerging shoots were each transferred to be propagated in a hydroponic environment with complete Hoagland solution (HiMedia Hoaglands No. 2 Basal Salt Mixture), keeping the light and temperature conditions consistent. Once the nettles grew to a size of four true leaves or more, they were separated into 100 mL Erlenmeyer flasks.

Further, the experiment involved four environmental conditions: a control group (no treatment), high light (HL), nitrogen deficiency (ND), and a combination of high light and nitrogen deficiency (HL+ND). Ten plants were kept under the previously established conditions while another ten stinging nettles were grown in either Hoagland solution without nitrogen (ND), high light conditions (HL) of 180-200 µmol/m2 s PPFD or both (HL+ND). At the end of a time period of two weeks, leaf color changes were observed and documented to assess the visual impact of the treatments.

For anthocyanin quantification, following steps were executed as previously described (Martin *et al*., 2002). First, approximately 25-100 mg of leaf tissue was collected, frozen at a temperature of -70 °C and then homogenized using a Precellys ribolyzer. The homogenization process was conducted at 6000 rpm for 45 seconds of on-time, followed by 60 seconds of off-time, repeated three times until the tissue was fully homogenized. Then, the resulting homogenate was centrifuged at 13,000 rpm for 1 minute, and the supernatant was discarded. To extract anthocyanins, 300 µL of 7% (v/v) HCl in methanol was added to each sample, and the samples were stored overnight at 4 °C. After an incubation period of 12–24 hours, 200 µL of deionized water and 500 µL of chloroform were added to the sample tubes. The tubes were mixed by inversion, followed by centrifugation at 13,000 rpm for 2 minutes. During this step, the tubes were kept on ice. After centrifugation, the upper aqueous phase (∼400 µL) was carefully removed and transferred to a fresh tube. Subsequently, 600 µL of 1% (v/v) HCl in methanol was added to each sample, and the samples were centrifuged again at 13,000 rpm for 2 minutes. Lastly, the absorbance of the supernatant was measured at 530 nm and 657 nm against the reagent blank, using a nanodrop spectrophotometer. Relative anthocyanin levels were determined using the following equation:

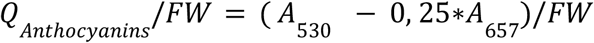

where *A530 and A657* represent the absorbance at 530 nm and 657 nm, respectively, and *FW* denotes the fresh weight (in grams) of the leaf tissue (Rabino & Mancinelli, 1986). For each biological replicate, the leaves were cut in half, and a separate measurement was taken from each half. The mean of these technical replicates was calculated to obtain a single value of *Qanthocyanins/FW* per biological replicate.

Then, the standard deviation (*σ*) was calculated using the following formula:

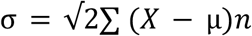

where *X* is each *Qanthocyanin/FW* value, *µ* represents the mean of all biological replicates, and *n* is the number of biological replicates. Afterwards the standard error was derived using this formula: 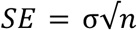

In this formula, *SE* stands for the standard error, *σ* is the population standard deviation and *n* is the number of elements. Further, the statistical significance of the differences between experimental groups was evaluated using a Kruskal-Wallis test for the analysis between the groups at day 0 and day 14. If a significant difference was revealed, a Dunn Post-Hoc test was performed. To determine the difference between day 0 and day 14 within each condition a Wilcoxon test was performed. Lastly, the data was visualized as a boxplot diagram.

## Availability of data and materials

All nanopore sequencing data have been submitted to the European Nucleotide Archive (ENA) and can be accessed through the project PRJEB63450. The *Urtica dioica* (XX-0-BONN-49798) genome sequence and corresponding annotation are available via bonndata (10.60507/FK2/1XHSZ3). Scripts used in this analysis are available via GitHub (https://github.com/PuckerLab/Urtica_dioica).

## List of abbreviations

SRE: Short Read Eliminator kit
ONT: Oxford Nanopore Technologies

## Declarations

### Ethics approval and consent to participate

Not applicable

### Consent for publication

Not applicable

### Authors’ contributions

KW and BP designed the project. KW and JAVSdV supervised and conducted the sequencing. KW and BP performed data analysis and wrote the initial draft of the manuscript. JAVSdV, LF, BG, MB, and MH contributed to the data generation and manuscript writing. MH contributed to the design of the figures. All authors have revised the manuscript and agreed to its submission.

## Acknowledgements

This work was supported by the de.NBI Cloud within the German Network for Bioinformatics Infrastructure (de.NBI) and ELIXIR-DE (Forschungszentrum Jülich and W-de.NBI-001, W-de.NBI-004, W-de.NBI-008, W-de.NBI-010, W-de.NBI-013, W-de.NBI-014, W-de.NBI-016, W-de.NBI-022). We also thank all students participating in our ‘Data Literacy in Genome Research’ course and members of the research group Plant Biotechnology and Bioinformatics for discussion and support. Elena Simeonidis extracted the RNA of *Urtica dioica* from different tissues for the RNA-seq experiments. Parts of this work are the result of the bachelor thesis written by Lena Fürstenberg. We are grateful for the excellent support provided by the team of the University of Bonn Botanic Gardens.

