## Supplementary figures and images for "Genome sequence of the medicinal plant *Urtica dioica* reveals the genetic basis of the flavonoid metabolism"

### Additional file B

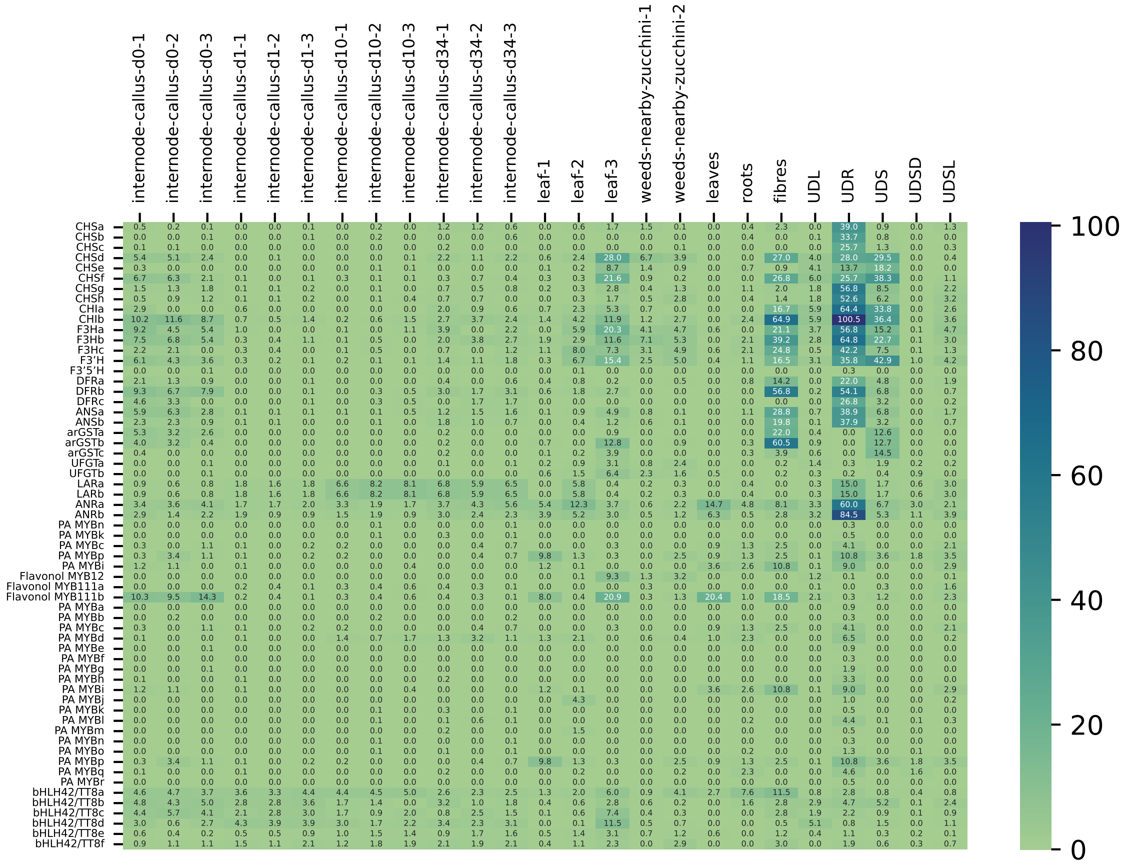

### Additional file E

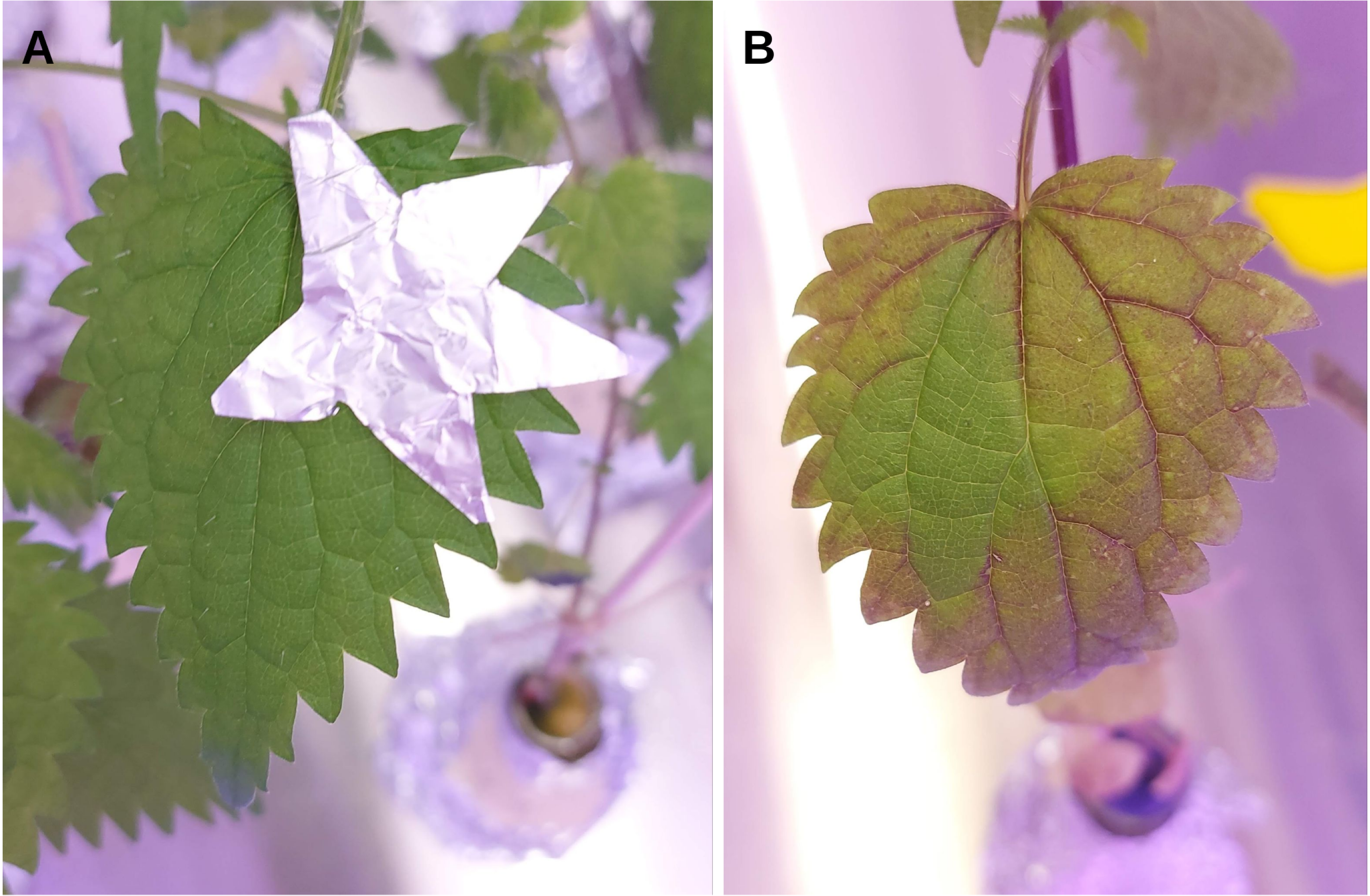
