## Additional file C for "Genome sequence of the medicinal plant *Urtica dioica* reveals the genetic basis of the flavonoid metabolism"

| Assembler | Shasta | Shasta | NextDenovo2 | NextDenovo2 | Verkko | Hifiasm |
|---|---|---|---|---|---|---|
| **Data** | R9 | R9 + R10 | R9 | R9 + R10 | R9 + R10 | R9 + R10 |
| **Number of contigs** | 7,562 | 938 | 1,464 | 1,455 | 2,329 | 639 |
| **Maximal contig length [bp]** | 9,134,718 | 30082981 | 11,616,341 | 15,011,099 | 34,769,096 | 59,598,013 |
| **Total number of bases [bp]** | 1,001,201,413 | 1,875,181,324 | 1,361,703,373 | 1,701,033,669 | 2,129,721,475 | 1,180,716,681 |
| **GC content [%]** | 41 | 41 | 41 | 41 | 41 | 42 |
| **N50 [bp]** | 327,375 | 6,089,958 | 1,424,355 | 1,972,031 | 6,831,518 | 32,829,641 |
| **N90 [bp]** | 68,438 | 1,437,265 | 467,741 | 559,481 | 511,358 | 5,919,930 |
| **BUSCO complete [%]** | 93.7 | 97.7 | 96.8 | 97.9 | 98.0 | 97.5 |
